# Mediator subunit MDT-15/MED15 and Nuclear Receptor HIZR-1/HNF4 cooperate to regulate toxic metal stress responses in *Caenorhabditis elegans*

**DOI:** 10.1101/565739

**Authors:** Naomi Shomer, Alexandre Zacharie Kadhim, Jennifer Margaret Grants, Xuanjin Cheng, Amy Fong-Yuk Poon, Michelle Ying Ya Lee, Forum Bhanshali, Anik Muhuri, Jung In Park, Dongyeop Lee, Seung-Jae V. Lee, Francis Christopher Lynn, Stefan Taubert

## Abstract

Zinc is essential for cellular functions as it is a catalytic and structural component of many proteins. In contrast, cadmium is not required in biological systems and is toxic. Zinc and cadmium levels are closely monitored and regulated as their excess causes cell stress. To maintain homeostasis, organisms induce metal detoxification gene programs through stress responsive transcriptional regulatory complexes. In *Caenorhabditis elegans,* the MDT-15 subunit of the evolutionarily conserved Mediator transcriptional coregulator is required to induce genes upon exposure to excess zinc and cadmium. However, the regulatory partners of MDT-15 in this response, its role in cellular and physiological stress adaptation, and the putative role mammalian for MED15 in the metal stress responses remain unknown. Here, we show that MDT-15 interacts physically and functionally with the Nuclear Hormone Receptor HIZR-1 to promote molecular, cellular, and organismal adaptation to excess metals. Using gain- and loss-of-function mutants and qPCR and reporter analysis, we find that *mdt-15* and *hizr-1* cooperate to induce zinc and cadmium responsive genes. Moreover, the two proteins interact physically in yeast-two-hybrid assays and this interaction is enhanced by the addition of zinc or cadmium, the former a known ligand of HIZR-1. Functionally, *mdt-15* and *hizr-1* mutants show defective storage of excess zinc in the gut, and at the organismal level, *mdt-15* mutants are hypersensitive to zinc- and cadmium-induced reductions in egg-laying. Lastly, mammalian MDT-15 orthologs bind genomic regulatory regions of metallothionein and zinc transporter genes in a metal-stimulated fashion, and human MED15 is required to induce a metallothionein gene in lung adenocarcinoma cells exposed to cadmium. Collectively, our data show that *mdt-15* and *hizr-1* cooperate to regulate metal detoxification and zinc storage and that this mechanism appears to be at least partially conserved in mammals.

## Introduction

In their habitats, biological organisms encounter many metals, including essential micronutrients such as iron, zinc, copper, and manganese, and toxic metals such as cadmium, mercury, lead, and arsenic. Zinc is an essential trace element that plays a crucial role in numerous cellular and physiological processes (1). It has a structural role in metabolic enzymes, growth factors, and transcriptional regulators such as zinc finger proteins, and is also an enzymatic cofactor and a signaling molecule (2,3). Accordingly, zinc is necessary for the function of approximately 10% of proteins in the human proteome and approximately 8% of proteins in the nematode worm *Caenorhabditis elegans* (4). In line with its requirement in diverse proteins, zinc deficiency causes a broad range of symptoms and dysfunctions in humans, such as skin and eye lesions, thymic atrophy, diarrhea, defective wound healing, and others (5,6). *Vice versa,* exposure to high doses of zinc is also detrimental, as it has toxic effects, causes cell stress, and alters physiological programs such as systemic growth, immune responses, and neuro-sensory and endocrine functions (5).

Unlike zinc, cadmium is a non-essential toxic metal encountered by biological organisms as a naturally occurring and industrial environmental contaminant. Cadmium has no known function in biological systems, and exposure causes intracellular damage along with the production of reactive oxygen species (7). In humans, cadmium exposure can result in respiratory disease, kidney damage, neurological disorders, and various types of cancers (7,8). Interestingly, zinc and cadmium share a similar electron configuration; thus, cadmium may substitute for zinc at the molecular level, for example as an enzyme cofactor, thus reducing or abrogating normal protein function (9).

Another consequence of the elemental similarity is that the biological responses to and the systemic detoxification of zinc and cadmium are similar (2,5,10,11). Key detoxification and homeostasis components include metal-sequestering proteins such as metallothioneins (MTs), which bind a wide range of metals including cadmium, lead, zinc, mercury, copper, and others. MTs appear to regulate the uptake, transport, and regulation of zinc in biological system, and also scavenge reactive oxygen species such superoxide (12). Other important detoxification and homeostasis components include transporters such as the cation diffusion facilitators (CDF; also known as zinc transporter (ZnT) or solute carrier 30 (SLC30) family proteins) that transport zinc into the cytoplasm, and the Zrt- and Irt-like proteins (ZIP; aka SLC39A family proteins) that transport zinc out of the cytoplasm (13,14).

To maintain homeostasis in the face of changing metal levels, transcriptional regulatory mechanisms adjust gene expression as needed. Metal-responsive transcription factor-1 (MTF-1) is a transcription factor that is evolutionarily conserved from insects to humans. It binds metal responsive elements (MREs) in the promoters of target genes (e.g. metallothioneins) and activates their expression when metals such as zinc are in excess (15–17). MTF-1 directly senses zinc with its six zinc fingers and with an acidic, metal-responsive transcriptional activation domain; related metals such as cadmium appear not to directly bind MTF-1 and instead are likely detected indirectly through the altered availability of zinc. Additional, unknown transcription factors and/or activation mechanism likely control gene expression in response to excess zinc in these organisms, as some genes are regulated independently of MTF-1 and MREs in such conditions (16,18). Recent studies in *C. elegans,* whose genome lacks a detectable MTF-1 ortholog despite the presence of MREs, revealed that the activation of genes in conditions of high zinc requires the High Zinc Activated (HZA) element and the HZA-binding Nuclear Hormone Receptor high-zinc–activated nuclear receptor 1 (HIZR-1; aka NHR-33) (19,20). HIZR-1 is a sequence homolog of mammalian Hepatocyte Nuclear Factor 4 (HNF4) (20,21), but whether HNF4 proteins regulate gene expression in response to high metal concentrations in mammals remains unknown.

To effectively and specifically activate gene expression, transcription factors require accessory proteins termed coregulators (22–24). One important coregulator is Mediator, a ~30 protein subunit complex that is conserved from yeast to human (25,26). Individual Mediator subunits selectively engage transcription factors and thus regulate specific developmental and physiological gene programs. For example, Mediator subunit MDT-15/Med15 and Mediator kinase cyclin dependent kinase 8 (CDK-8) are required for many stress and adaptive responses across species (26–31). In the context of heavy metal responsive transcription, *Drosophila melanogaster* MTF-1 requires Mediator for gene activation via MREs in response to excess metals (17,32). In *C. elegans,* we showed that Mediator subunit *mdt-15* is required for the induction of zinc and cadmium responsive genes (33). Others found that zinc-dependent activation via the HZA element required *mdt-15* at one extrachromosomal promoter reporter (19). As MDT-15 is a known coregulator of HNF4-like NHRs in *C. elegans* (34–37), this suggests that MDT-15 may cooperate with the HZA-binding HIZR-1 to adapt gene expression in response to high zinc in *C. elegans.*

Here, we tested whether MDT-15 and HIZR-1 cooperate to regulate zinc and cadmium responsive transcription in *C. elegans* and whether MED15, the mammalian ortholog of MDT-15, also participates in heavy metal stress responses. Because the Mediator kinase CDK-8 regulates transcriptional stress responses (30,31), we also assessed the function of *C. elegans cdk-8* in metal responsive transcription. Using genetic, molecular, cytological, and functional assays, we find that MDT-15 and HIZR-1 interact physically and functionally in zinc and cadmium stress responses, and that mammalian MED15 is recruited to metal responsive genes and required for metal-induced gene expression.

## Results

### *mdt-15* and *cdk-8* are necessary for cadmium- and zinc-induced gene activation

We previously showed that *mdt-15* is required to induce the mRNA levels of several genes in response to high levels of zinc and cadmium (33). In line with this finding, genes downregulated in *mdt-15(tm2182)* hypomorph mutants showed significant overlap with cadmium-induced genes (Fig 1A). Similarly, cadmium-induced genes overlap significantly with genes downregulated in *cdk-8 (tm1238)* null mutants (Fig 1B), suggesting that *cdk-8* plays a similar role as *mdt-15.* To test the requirement of *cdk-8* in cadmium detoxification and to assess its role in the zinc response, we used real-time quantitative PCR (qPCR) to measure the mRNA levels of metallothioneins (*mtl-1, mtl-2*), zinc transporters (ZnT) implicated in cadmium and zinc detoxification (*ttm-1, cdf-2*), and the cadmium responsive gene *cdr-1* (10,20,33). Comparing wild-type worms and *cdk-8(tm1238)* mutants, we observed a significant decrease of *cdr-1* mRNA levels upon exposure to cadmium and of *mtl-2* mRNA levels upon exposure to zinc; other metal responsive genes showed a similar trend but were not significantly altered (Fig 1C-D). Thus, *cdk-8* is required for maximal gene expression in response to zinc or cadmium, but this requirement is less prevalent and substantial than the one we previously observed for *mdt-15* (33).

**Figure 1:**
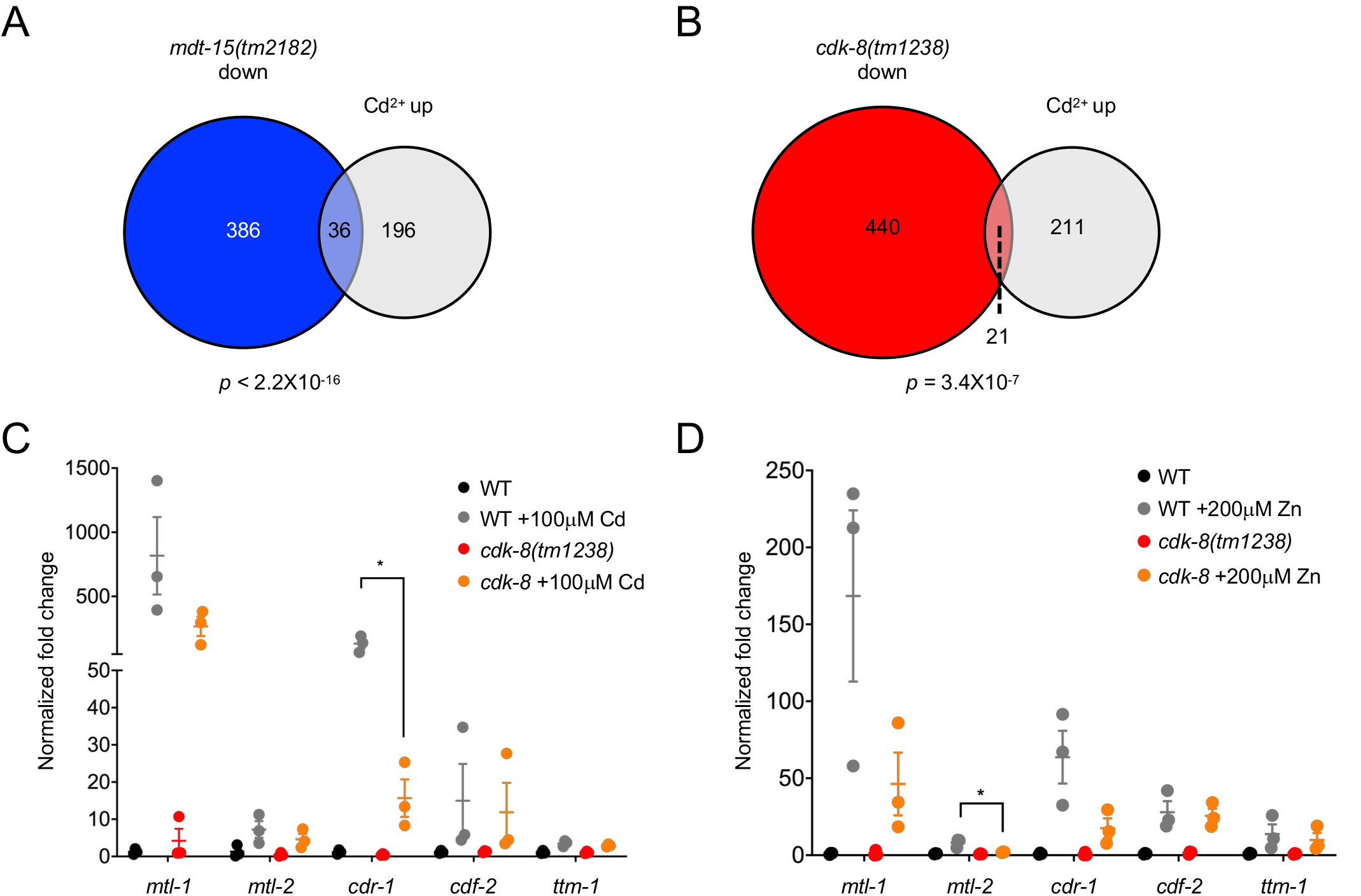
Mediator subunits *mdt-15* and *cdk-8* are required for heavy metal stress responses. **[A]** The Venn diagram depicts the overlap between genes downregulated in the *mdt-15(tm2182)* mutant and genes induced by cadmium (from (62)). Statistical significance was assessed by Fisher’s exact test. **[B]** The Venn diagram depicts the overlap between genes downregulated in the *cdk-8(tm1238)* mutant and genes induced by cadmium. Statistical significance was assessed by Fisher’s exact test. **[C-D]** qPCR analysis of zinc and cadmium-responsive genes in wild-type (WT) and *cdk-8(tm1238)* mutant worms grown on 100μM cadmium for 4 hours [C] or on 200μM zinc for 16 hours [D]. Graphs show fold induction, normalized to the average of unsupplemented WT mRNA levels. Error bars: SEM (*n*=3). Statistical analysis: * *p* < 0.05, unpaired t-test comparing WT supplemented worms to mutant supplemented worms.

### *hizr-1* and *elt-2* are required to induce the *cdr-1* promoter

To delineate the mechanism of MDT-15 and CDK-8 driven, cadmium and zinc responsive transcription, we generated a transcriptional *cdr-1p::gfp* reporter, encompassing 2.8 kb of the *cdr-1* promoter (Fig 2A). We chose *cdr-1* as a model because it is highly cadmium and zinc responsive and requires *mdt-15* and *cdk-8* for activation (10,33,38), suggesting that it might be a good tool to identify DNA regulatory elements and cognate transcription factors that cooperate with these Mediator subunits. As expected (10,33,38), we observed weak basal expression of this reporter, but substantial induction of fluorescence by 200μM zinc and 100μM cadmium (Fig S1A-B); expression was primarily localized to the intestine (Fig S1A-B), as expected (10,38). Knockdown of *mdt-15* by feeding RNA interference (RNAi) caused a significant decrease in basal *cdr-1p::gfp* fluorescence and abrogated fluorescence induction by cadmium and zinc (Fig 2B). Similarly, *cdk-8(tm1238); cdr-1p::gfp* worms showed a significant decrease in basal and metal-induced fluorescence (Fig 2B).

**Figure 2:**
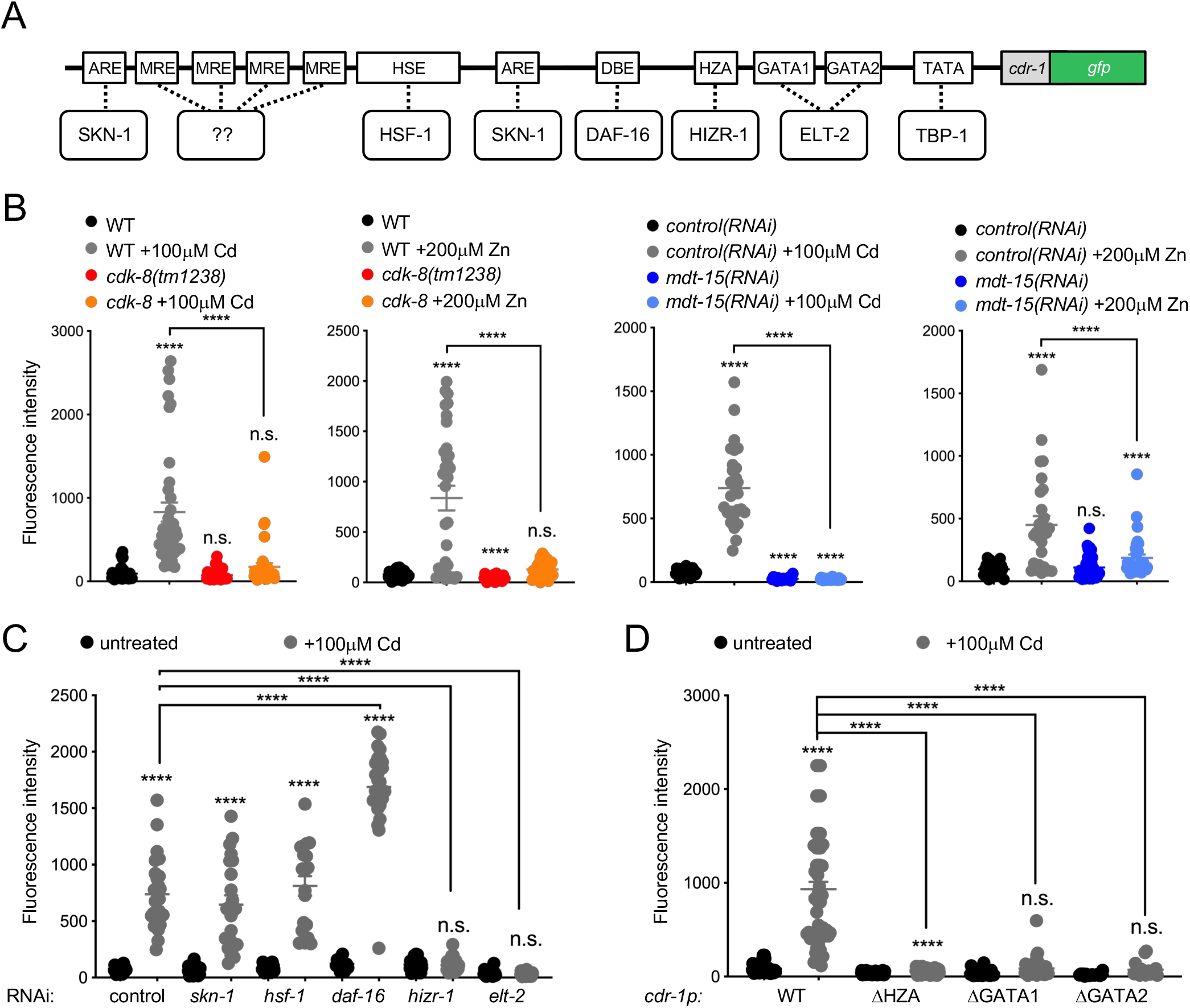
The TFs *hizr-1* and *elt-2* and their cognate promoter elements are required for heavy metal stress responses. **[A]** Illustration of the *cdr-1p::gfp* reporter and putative regulatory DNA elements in the *cdr-1* promoter; note, the diagram is not to scale. For details, see text. **[B]** Graphs show average fluorescence intensity (arbitrary units, A.U.) of worms bearing the *cdr-1p::gfp* transcriptional reporter in WT and *cdk-8(tm1238)* backgrounds or fed either control or *mdt-15* RNAi and exposed for four hours to 0μM or 100μM cadmium or 0μM or 200μM zinc, as indicated. Error bars: SEM (*n* > 19 worms per group). Statistical analysis: **** *p* < 0.0001, n.s.= not significant, Two-way ANOVA, multiple comparisons, Tukey correction; comparisons are to WT or *control(RNAi)* unless indicated. **[C]** Graph shows average fluorescence intensity (arbitrary units, A.U.) of worms bearing the *cdr-1p::gfp* transcriptional reporter before and after a 0μM or 100μM cadmium exposure (four hours), and fed either control or transcription factor RNAi, as indicated. Error bars: SEM (n > 19 worms per group). Statistical analysis: \*\*\*\**p* < 0.0001, n.s.= not significant, Two-way ANOVA, multiple comparisons, Tukey correction; comparisons are to *control(RNAi)* unless indicated. **[D]** Graph shows average fluorescence intensity (arbitrary units, A.U.) of worms bearing the following *cdr-1p::gfp* reporter variants: wild-type, WT; mutated HZA element, *cdr-1PΔHZA;* mutated GATA elements, *cdr-1PΔGATA1* or *cdr-1PΔGATA2.* All mutants were assessed with 0μM or 100μM cadmium (four hours). Error bars: SEM (n > 19 worms per group). Statistical analysis: **** *p* < 0.0001, n.s.= not significant, Two-way ANOVA, multiple comparisons, Tukey correction; comparisons are to WT unless indicated.

To identify transcription factors that cooperate with *cdk-8* and *mdt-15* to regulate cadmium and zinc responsive transcription, we searched for DNA regulatory elements in the *cdr-1* promoter. We identified candidate elements recognized by SKN-1/Nrf2 (antioxidant response element, ARE), HSF-1 (heat shock response element, HSE), DAF-16/FOXO (DAF-16 binding element, DBE), ELT-2 (GATA element), and HIZR-1 (HZA element), as well as four MREs, which no *C. elegans* transcription factor is yet known to bind (Fig 2A). RNAi analysis revealed that *skn-1, hsf-1,* and *daf-16* are not required for *cdr-1* induction by cadmium; *daf-16* depletion actually hyper-induced the *cdr-1p::gfp* promoter (Fig 2C). In contrast, knocking down *elt-2* or *hizr-1* abrogated fluorescence induction (Fig 2C). As *elt-2* is required for intestinal development (39), we examined post-developmental *elt-2* knockdown, which also caused abrogation of basal and cadmium-induced *cdr-1p::gfp* expression (Fig S1C). Thus, *elt-2* is required at the *cdr-1* promoter independently of its role in development.

We confirmed the requirements for *elt-2* and *hizr-1* by site-directed mutagenesis of their cognate DNA elements. We generated substitution mutations in the HZA (*mutHZA*) or GATA sites (*mutGATA1* and *mutGATA2)* of the *cdr-1p::gfp* reporter (Fig 2A). Each mutation individually caused a significant decrease in cadmium-induced promoter activity compared to the wild-type *cdr-1p::gfp* reporter; mutations in the HZA elements also caused a significant decrease in the basal activity of the *cdr-1p::gfp* reporter (Fig 2D). Collectively, these data show that Mediator subunits MDT-15 and CDK-8 and the transcription factors ELT-2 and HIZR-1 are required to control expression from the cadmium/zinc-inducible *cdr-1* promoter.

### *mdt-15* and *hizr-1* function is co-dependent

Based on the above data, we hypothesized that MDT-15 and/or CDK-8 might interact functionally and physically with HIZR-1 and/or ELT-2 to activate metal-induced transcription. To examine a putative functional relationship between MDT-15 and HIZR-1, we studied the *hizr-1(am285)* D270N gain-of-function (gf) mutant that induces zinc responsive genes even in the absence of zinc (20). In line with published data (20), qPCR analysis revealed induction of *cdr-1, mtl-1, mtl-2,* and *cdf-2* in *hizr-1(am285)* mutants grown on control RNAi; importantly, *mdt-15* RNAi significantly reduced or abrogated these inductions (Fig 3A). Next, we studied the *mdt-15(et14)* P117L gain-of-function (gf) mutation that induces MDT-15 regulated lipid metabolism genes (40). Because this mutation is closely linked to a *paqr-1(3410)* loss-of-function mutation, which it suppresses (40), we used the *mdt-15(yh8)* strain, which was generated by CRISPR and carries P117L alone (41). The *mdt-15(yh8)* gf mutation was sufficient to induce *mtl-1* and *mtl-2* (Fig 3B); RNAi experiments revealed that *mtl-2* induction required *hizr-1,* with a similar trend for *mtl-1* (Fig 3B). In contrast, *nhr-49,* which cooperates with MDT-15 to activate lipid metabolism and stress response genes (34,36), was not required to induce metal-responsive genes in the *mdt-15(yh8)* mutant (Fig 3B). We conclude that MDT-15 and HIZR-1 interact specifically to induce metal responsive genes *in vivo.*

**Figure 3:**
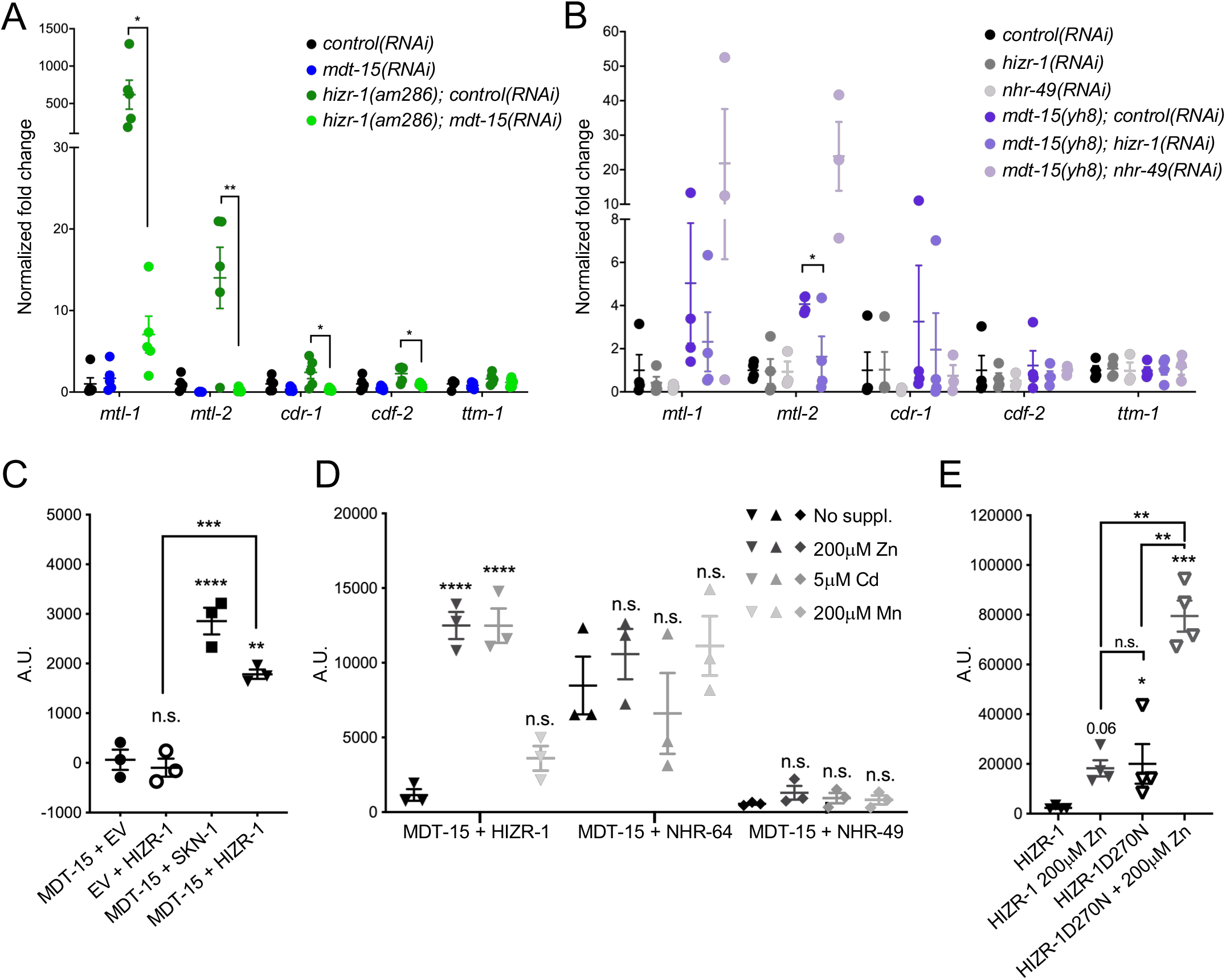
MDT-15 and HIZR-1 are co-dependent for metal responsive gene induction and physically bind in zinc-enhanced fashion. **[A-B]** qPCR analysis of zinc and cadmium-responsive genes. The graphs show fold-inductions of mRNAs normalized to the average of wild-type *control(RNAi)* worms. Error bars: SEM. Statistical analysis: \**p* < 0.05, \*\**p* < 0.01, unpaired Student’s t-test for indicated comparisons. [A] Comparison of wild-type and *hizr-1(am285)* worms grown on control RNAi or *mdt-15* RNAi (n=5). [B] Comparison of wild-type and *mdt-15(yh8)* worms grown on control RNAi, *hizr-1* RNAi, or *nhr-49* RNAi (n= 3). **[C-E]** Protein-protein interaction analysis using the Y2H system. Graphs show average interaction strength (arbitrary units, A.U.). Error bars: SEM. [C] Interaction between MDT-15 and HIZR-1, with Empty Vector (EV)-HIZR-1 and MDT-15-EV as negative controls, and MDT-15–SKN-1c as a positive control (n=3). Statistical analysis: *** p < 0.001, **** p < 0.0001, One-way ANOVA, multiple comparisons, Dunnett correction. All comparisons for “MDT-15 + EV” unless indicated. [D] Interaction strength between MDT-15 and HIZR-1; MDT-15 and NHR-64; and MDT-15 and NHR-49, all with no treatment, or treated with 200μM zinc, 5μM cadmium, or 200μM manganese (n=3). Statistical analysis: **** p < 0.0001, Two-way ANOVA, multiple comparisons, Dunnett correction. All comparisons to the pertinent “no supplement” control. [E] Interaction between HIZR-1 and MDT-15 or HIZR-1-D270N and MDT-15 with and without zinc treatment (n=4). Statistical analysis: ** p < 0.01, *** p < 0.001, One-way ANOVA, multiple comparisons, Dunnett correction. All comparisons to “HIZR-1” unless indicated.

### MDT-15 physically interacts with HIZR-1 in zinc-enhanced fashion

The above data suggest that MDT-15 might interact physically with HIZR-1, as shown for other *C. elegans* HNF4-like NHRs (34,35). To test whether HIZR-1 binds MDT-15, we used the yeast-two-hybrid (Y2H) system. First, we examined the interaction between a full length HIZR-1 prey and a MDT-15-ΔCT bait (aa 1-600); full-length MDT-15 (aa 1-780) autoactivates and cannot be used as bait in Y2H assays (see (42) for details). In these assays, MDT-15 and HIZR-1 showed a statistically significant interaction that was similar in strength to the interaction of the MDT-15-ΔCT bait with a SKN-1c prey (positive control (42); Fig 3C).

The HIZR-1 ligand binding domain (LBD) binds zinc in the micromolar range, suggesting that zinc is a *bona fide* ligand for HIZR-1 (20). We hypothesized that the addition of zinc might enhance the interaction of HIZR-1 with MDT-15, as shown for other many other ligand-stimulated NHR-coregulator interactions. Indeed, addition of zinc enhanced the interaction between MDT-15 and HIZR-1, with significant effects at low micromolar concentrations (Figs 3D, S2A). As *mdt-15* also contributes to cadmium-induced gene expression, we tested whether this metal also enhanced MDT-15 binding to HIZR-1 and found that this was the case (Fig 3D). In contrast, manganese did not enhance the binding of MDT-15 with HIZR-1 (Fig 3D).

NHRs contain a zinc-finger DNA binding domain (DBD), suggesting that zinc-stimulated binding to a coregulator such as MDT-15 might be a common feature of NHRs. To examine the specificity of zinc- and cadmium-stimulated MDT-15–NHR interaction, we examined whether these metals alter the binding of two other known MDT-15 binding partners, NHR-64 and NHR-49 (34). We found that NHR-64 interacts with MDT-15 as strongly as HIZR-1 does in the presence of zinc or cadmium, while NHR-49 binding to MDT-15 mimics the interaction of HIZR-1 and MDT-15 in the absence of zinc; importantly, neither NHR-64 nor NHR-49 binding to MDT-15 was altered by supplementation with zinc, cadmium, or manganese at eth concentrations that significantly enhanced binding of HIZR-1 to MDT-15 (Fig 3D). We conclude that metal-stimulation of MDT-15 interaction is not a general feature of NHRs but specific to HIZR-1.

### Binding determinants in MDT-15 and HIZR-1

MDT-15 contains an N-terminal KIX-domain that binds several NHRs and the lipogenic transcription factor SBP-1 (34,35,43). Thus, we hypothesized that MDT-15 might physically bind HIZR-1 through the KIX-domain (aa 1-124; Fig S2B). To test this hypothesis, we assayed binding of HIZR-1 to an MDT-15-KIX-domain and to an MDT-15ΔCT variant lacking the KIX-domain (MDT-15ΔKIXΔCT; aa 125-600). The binding of HIZR-1 to the MDT-15-KIX-domain was similar to the binding to MDT-15ΔCT; in contrast, MDT-15ΔKIXΔCT was unable to bind HIZR-1 (Fig S2C). This indicates that the KIX-domain is necessary and sufficient for the HIZR-1–MDT-15 interaction.

The *hizr-1(am285)* gf mutation is an aspartate 270 to asparagine (D270N) substitution that results in increased nuclear localization and constitutive activation of zinc responsive genes even in the absence of zinc (20). Sequence comparison revealed that D270 is conserved in the HIZR-1 orthologs of three other species in the *Caenorhabditis* genus (Fig S2D), suggesting that it may be functionally important. We hypothesized that the D270N mutation might affect MDT-15 binding. To test this, we performed Y2H assays with a HIZR-1-D270N prey generated by site-directed mutagenesis. HIZR-1-D270N interacted more strongly with MDT-15 than did HIZR-1-WT, resembling in strength the WT HIZR-1–MDT-15 interaction in the presence of zinc (Fig 3E). Nevertheless, supplemental zinc further enhanced this interaction (Fig 3E), suggesting that, D270 does not mimic the effects of zinc-stimulated binding.

### *mdt-15(tm2182)* mutants display zinc storage defects and are hypersensitive to zinc and cadmium

To test whether the defects of *mdt-15(tm2182)* mutants in metal responsive transcription have functional consequences, we studied two phenotypes: cellular zinc storage and egg-laying. At the cellular level, gut granules store zinc when it is present in excess, thus protecting cells; *vice versa,* they replenish zinc in situations of zinc deficiency (44). To study zinc storage in gut granules in wild-type and mutant worms, we used the zinc-specific fluorescent dye FluoZin-3 (44). We observed little difference between wild-type, *mdt-15(tm2182), cdk-8(tm1238*), and *hizr-1(am286)* mutants in “normal” conditions, i.e. without zinc supplementation (Fig S3A-C). Wild-type worms supplemented with 200mM zinc showed bigger granules with a stronger FluoZin-3 signal (Fig 4A), as reported (44). Strikingly, *mdt-15(tm2182)* mutants displayed significantly fewer gut granules in high zinc conditions than wild-type worms and *cdk-8(tm1238)* mutants (Fig 4A-B). As HIZR-1 interacts with MDT-15 in zinc-stimulated fashion, we also studied *hizr-1(am286)* null mutants and found that they also have less granules than wild-type worms in conditions of excess zinc (Fig 4C).

**Figure 4:**
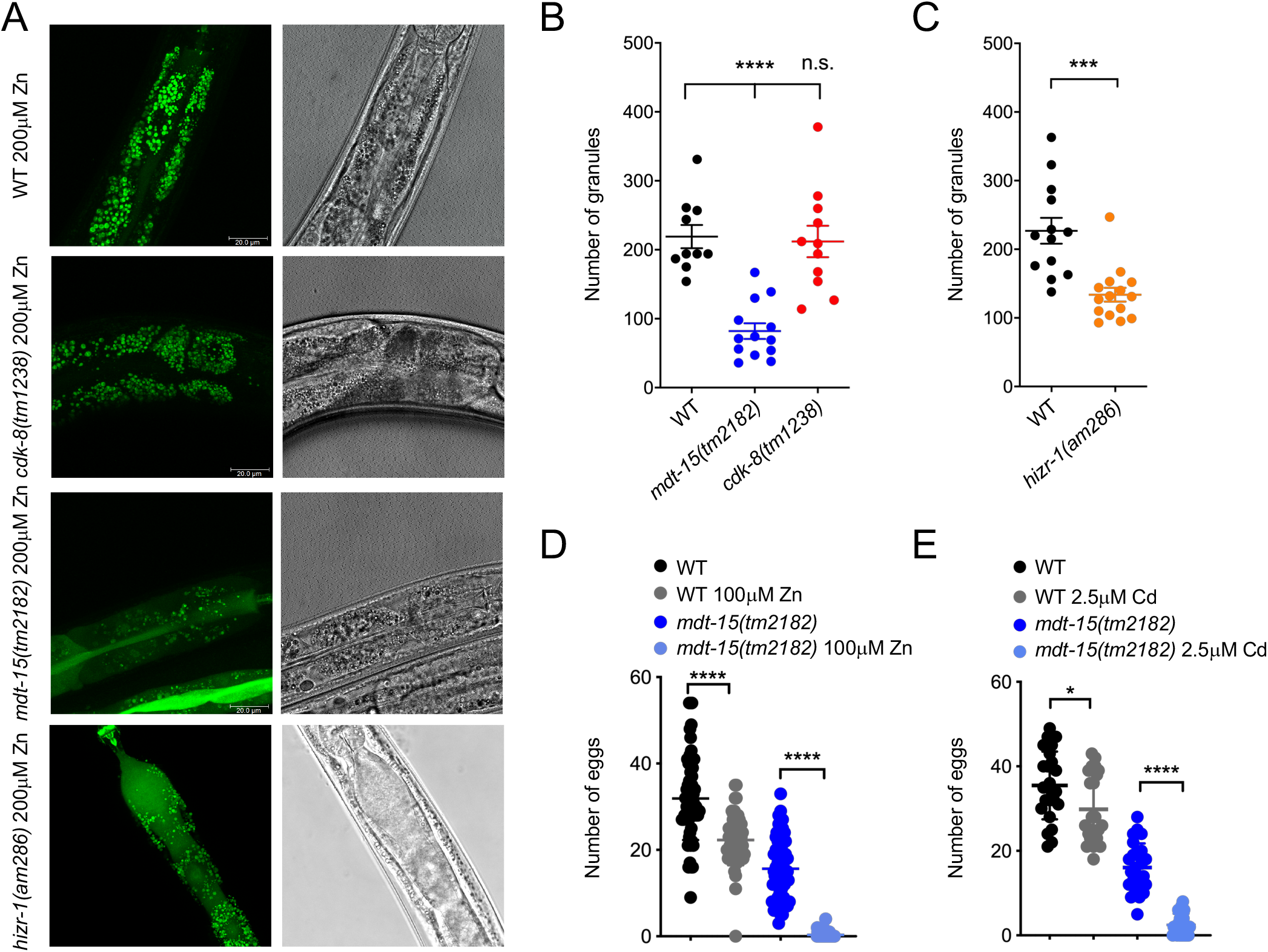
*mdt-15(tm2182)* mutants have a zinc storage defect and are sensitive to excess heavy metals. **[A]** Representative fluorescence images of worms stained with FluoZin-3, showing zinc accumulation in gut granules in wild-type, *mdt-15(tm2182), cdk-8(tm1238*), and *hizr-1(am286)* worms grown with 200μM zinc supplementation. **[B]** Quantification of the number of gut granules in wild-type (WT), *mdt-15(tm2182*), and *cdk-8(tm1238)* worms; every dot represents an individual worm. Error bars: SEM. *n* > 10 worms per genotype. Statistical analysis: **** p < 0.0001, One-way ANOVA, multiple comparisons, Dunnett correction, compared to “WT”. **[C]** Quantification of the number of gut granules in WT and *hizr-1(am286)* worms; every dot represents an individual worm. Error bars: SEM. *n* > 13 worms per genotype. Statistical analysis: *** p < 0.001, One-way ANOVA, multiple comparisons, Dunnett correction, compared to “WT”. **[D]** Graphs show the average number of eggs laid over a 24-hour period by WT worms and *mdt-15(tm2182)* mutants grown on 0μM and 100μM zinc (n > 51 worms per condition). Error bars: SEM; statistical analysis: **** *p* < 0.0001 Two-way ANOVA, multiple comparisons; Tukey correction. **[E]** Graphs show the average number of eggs laid over a 24-hour period by WT worms and *mdt-15(tm2182)* mutants grown on 0μM and 2.5μM cadmium (n > 23 worms per condition). Error bars: SEM; statistical analysis: \*\*\*\**p* < 0.0001 Two-way ANOVA, multiple comparisons; Tukey correction.

*mdt-15(tm2182)* mutants show reduced expression of several fatty acid metabolism enzymes, especially fatty acid desaturases such as *fat-6/stearoyl-CoA desaturase*. This causes defects in cell membrane fatty acid desaturation and directly underlies numerous phenotypes caused by *mdt-15* deficiency, including slow growth, reduced body size, and short life span (34,41,43,45,46). Altered membrane lipids in *mdt-15(tm2182)* mutants could cause organelle dysfunction and conceivably affect zinc storage. Thus, we tested whether *fat-6/stearoyl-CoA desaturase* depletion by RNAi causes defects in zinc storage (note that *fat-6* depletion also deletes the highly homologous gene *fat-7* (47)). In contrast to *mdt-15(tm2182)* mutants, however, *fat-6(RNAi)* worms did not show any overt defects in zinc storage in high zinc conditions (Fig S3D-E). This suggests that the zinc storage defects observed in the *mdt-15(tm2182)* mutants are not due to altered membrane composition and function.

We also assessed the effect of excess zinc and cadmium on an organismal phenotype, egg-laying (Fig 4D). We found that 100μM zinc decreased the number of eggs laid by wild-type worms by approximately 30 percent; in contrast, the same concentration of zinc almost completely abolished egg-laying in *mdt-15(tm2182)* mutants, indicating that this mutant is hyper-sensitive to zinc. Similarly, 2.5μM cadmium reduced the number of eggs laid by wild-type worms by approximately 20 percent, but almost completely abrogated it in *mdt-15(tm2182)* mutants (Fig 4E), indicating that this mutant is hyper-sensitive to cadmium. Thus, *mdt-15(tm2182)* mutants are less able than wild-type to resist excess heavy metals at an organismal level.

### Mammalian MED15 may also regulate metal-induced transcription

To test whether *mdt-15*’s role in regulating metal-responsive transcription is conserved in mammals we studied the role its human and mouse orthologs, MED15 and Med15 (48), in two cell lines. First, we studied A549 human epithelial lung adenocarcinoma cells, because cadmium increases lung cancer risk and induces a heavy metal stress response in this cell line (49–51). We depleted MED15 in A549 cells by transfecting small interfering RNAs (siRNAs) targeting MED15 (vs. scrambled control), and then exposed the transfected cells to 5μM cadmium for 4 hours. Using qPCR, we found that cadmium induced the metallothioneins MT1X and MT2A, orthologues of *C. elegans mtl-1* and −2, respectively (Fig 5A). Notably, while MT1X was unaffected, MT2A induction was completely blocked by MED15 depletion (Fig 5A), suggesting that MED15 is required to induce at least this one gene in response to heavy metal exposure. To test whether MED15 binds to the promoter of MT2A, we performed chromatin immunoprecipitation followed by qPCR (ChIP-qPCR), assessing MED15 occupancy before and after the addition of 5μM cadmium to the extracellular media. We found that MED15 was recruited preferentially to the promoter of MT2A after addition of 5μM cadmium (Fig 5B).

**Figure 5.**
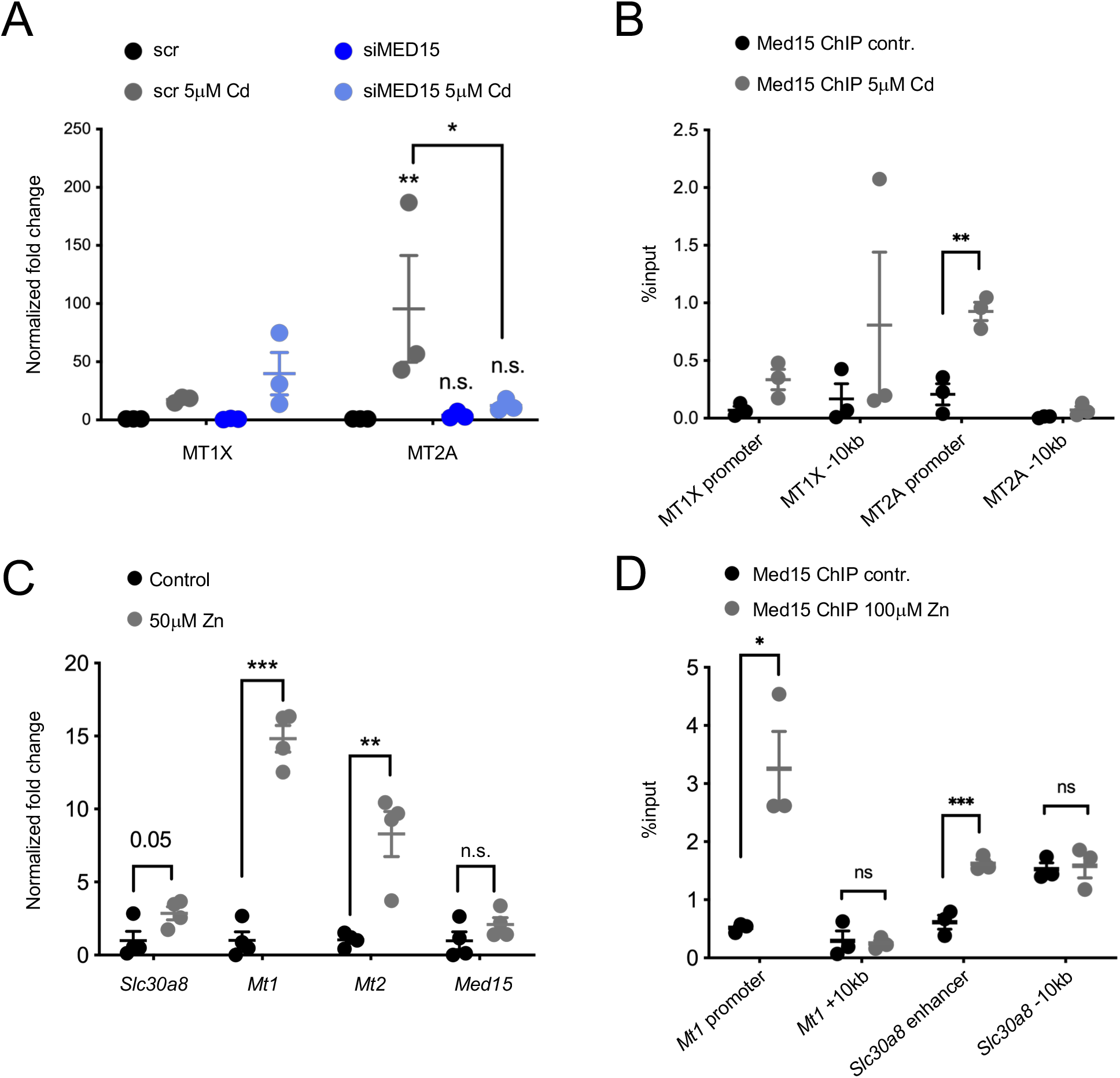
MED15 binds and regulates metal responsive genes in mammalian cells. **[A]** qPCR analysis of cadmium inducible genes in control (scr = scrambled siRNA) and MED15 siRNA (siMED15) treated A549 cells exposed to 5μM cadmium for four hours. MT1X was not affected in statistically significant fashion. For MT2A, all comparisons are *vs.* the untreated scr samples except where indicated. Error bars: SEM; statistical analysis: * *p* < 0.05, \*\**p* < 0.01 Two-way ANOVA, multiple comparisons, Tukey correction (n=3). **[B]** The graph shows relative MED15 occupancy at the MT1X and MT2A promoters and nearby control regions in A549 cells, as determined by ChIP-qPCR before and after the addition of 5μM cadmium. Error bars: SEM; statistical analysis: ** *p* < 0.01 Student’s T-test, multiple comparisons, Holm-Sidak correction (n=3). **[C]** The graph shows qPCR analysis of MIN6 cells treated without and with 50μM zinc. Error bars: SEM; statistical analysis: ** *p* < 0.01, *\*\*\*p* < 0.001 Student’s T-test, multiple comparisons, Holm-Sidak correction (n=4). **[D]** The graph shows relative Med15 occupancy at the *Mt1* promoter and Slc30a8 enhancer and nearby control regions in MIN6 cells as determined by ChIP-qPCR, before and after the addition of 100μM Zn. Error bars: SEM; statistical analysis: *\*p* < 0.05, *\*\*\*p* < 0.01 Student’s T-test, multiple comparisons, Holm-Sidak correction (n=3).

We also studied the MIN6 mouse insulinoma cell line because the insulin-secreting β-cells of the mammalian pancreas require zinc for insulin crystallization and contain among the highest levels of zinc in the body (14,52,53). ZnT8/SLC30A8, the mouse ortholog of the MDT-15 and HIZR-1 regulated zinc transporter CDF-2 (44), is expressed highly in the in α- and β-cells of the endocrine pancreas (14). To test whether the expression of mouse *Slc30a8* and metallothionein genes are induced by excess zinc, we exposed MIN6 cells to 50μM zinc for 24 hours and assessed expression by qPCR. We observed that *Slc30a8, Mt1,* and *Mt2* are zinc responsive, whereas *Med15* expression did not increase (Fig 5C). To test whether Med15 directly binds to the promoters of *Slc30a8* and *Mt1,* we performed chromatin immunoprecipitation followed by qPCR (ChIP-qPCR), assessing Med15 occupancy before and after the addition of excess zinc to the extracellular media. We found that Med15 was recruited to the promoters of both *Slc30a8* and *Mt1* in excess zinc (Fig 5D). Collectively, these experiments suggest that mammalian MED15 proteins directly bind the promoters of metal stress responsive genes and are required for the induction of at least one cadmium responsive gene in lung adenocarcinoma cells.

## Discussion

Organisms encounter both essential and toxic metals in their habitant and must allow adequate uptake of necessary micronutrients while secreting/sequestering excess amounts of micronutrients and toxic metals. *C. elegans* lacks the MTF-1 protein that regulates metal responsive transcription in many animals. Instead, it utilizes the Nuclear Hormone Receptor HIZR-1 to control the response to excess zinc (20). Here, we show that the *C. elegans* Mediator subunit MDT-15 interacts physically and functionally with HIZR-1, that loss of *mdt-15* alters storage of excess zinc *in vivo,* and that *mdt-15* mutants are hypersensitive to zinc and cadmium. Moreover, mammalian MED15 is recruited to regulatory elements of metal-responsive genes in metal-stimulated fashion, and MED15 depletion blocks the induction of a cadmium responsive gene in lung adenocarcinoma cells. Thus, the HZA element, HIZR-1 transcription factor, and MDT-15 coregulator compose a regulatory mechanism that adapts gene expression in response to excess zinc and cadmium to protect the host organism (Fig 6). This regulatory mechanism represents a new partnership between a *C. elegans* NHR and the coregulator MDT-15 that regulates one particular adaptive response, that to excess heavy metals. Moreover, this metal stress response mechanism appears to be at least partly conserved in mammalian cells (Fig 6).

**Figure 6.**
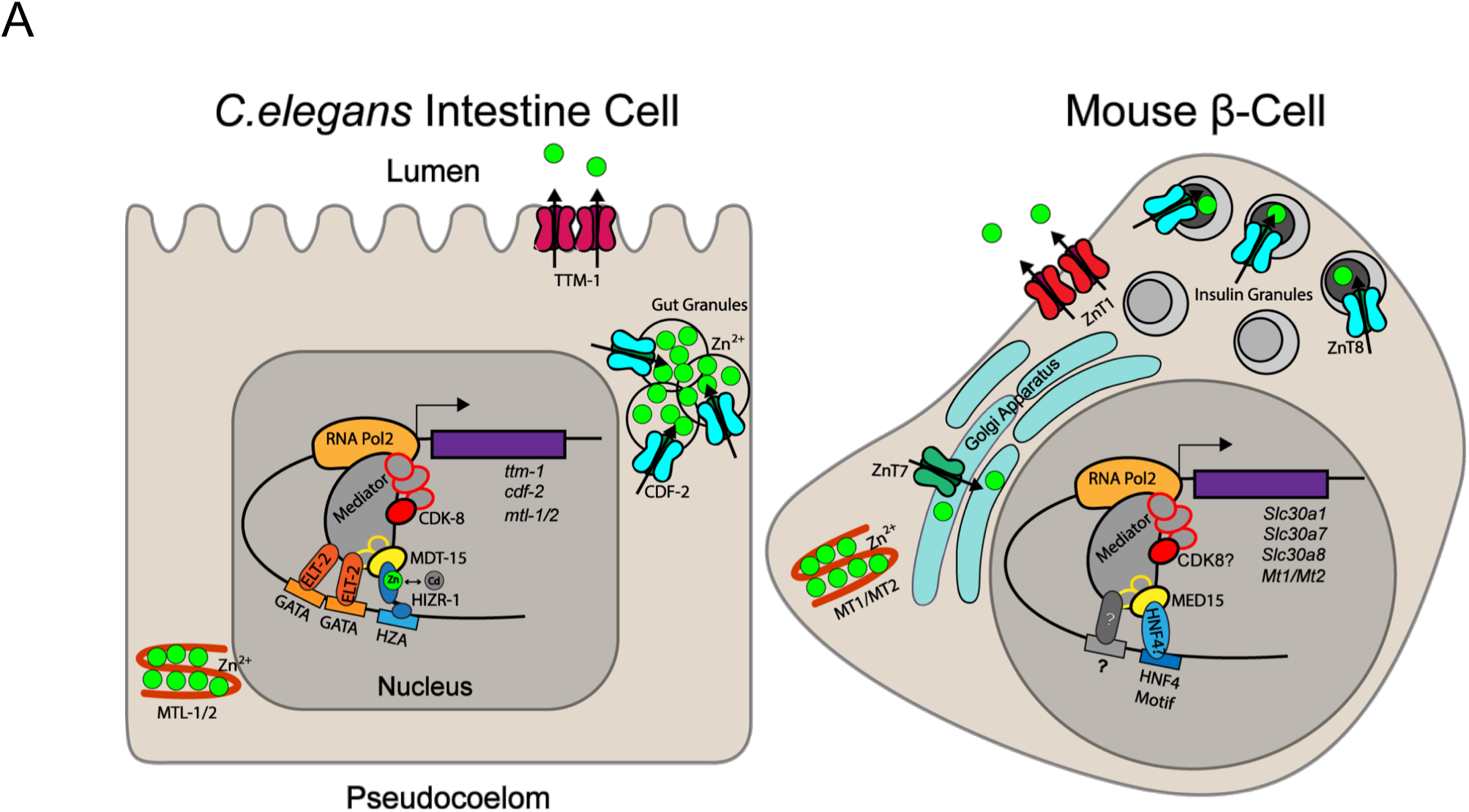
Model of MDT-15 and MED15 driven gene regulation in response to high metal levels. **[A]** In the *C. elegans* intestinal cells, MDT-15 cooperates with HIZR-1 bound to the HZA element to induce the metal sequestering metallothioneins (*mtl-1* and −2) and the transporter *cdf-2,* which shuttles zinc into storage granules; excess cadmium is likely detoxified through similar mechanisms. ELT-2 may also contact Mediator to regulate metal responsive transcription. **[B]** In the mouse pancreatic β-cells, Med15 regulates the expression of genes such as Slc30a8/ZnT1 in response to elevated zinc levels, possibly via HNF4, or other, Mediator contacting proteins.

### MDT-15 is a coregulator of HIZR-1

Because *C. elegans* appears to lack an MTF-1 ortholog, it has been unclear how it regulates gene expression in response to changing levels of various metals. We previously showed that the Mediator subunit *mdt-15* is essential for both zinc and cadmium activated transcription (33), and the Kornfeld laboratory identified the HZA element and its cognate TF HIZR-1 as regulators of zinc-inducible transcription (19,20). This raised the hypothesis that these three components might all cooperate mechanistically; alternatively, they might act in separate molecular pathways. Here, we provide multiple lines of evidence that HIZR-1 and MDT-15 cooperate mechanistically to induce gene expression in excess zinc and cadmium. First, *mdt-15, hizr-1,* and the HZA element are required to activate the *cdr-1p::gfp* reporter, i.e. they phenocopy (Fig 2; see also (19,20)). Second, MDT-15 and HIZR-1 physically interact in the yeast-two-hybrid system; notably, this binding can be stimulated by a known ligand of HIZR-1 (zinc; (20)) and requires a known NHR binding domain of MDT-15 (the KIX-domain; (34,35); Fig 3). Third, genetic gain of *hizr-1* or *mdt-15* function confers metal homeostasis gene activation that requires the reciprocal partner (Fig 3). Fourth, loss of *mdt-15* and *hizr-1* causes similar defects in zinc storage and renders worms hypersensitive to zinc (Fig 4 and (20)). Although we cannot exclude that the GATA factor ELT-2, which also activates zinc responsive genes including *cdr-1* (Fig 2 and (19)), may interact with MDT-15, we have not observed an interaction between ELT-2 and MDT-15 in our yeast-two-hybrid assays. Based on our new and published data, we propose a model whereby elevated levels of zinc or cadmium promote the formation of a HIZR-1–MDT-15 regulatory complex, which acts through the HZA element to induce the expression of genes required for metal homeostasis (Fig 6).

### MDT-15 binds HIZR-1 in metal-stimulated fashion

Interestingly, zinc or cadmium supplementation enhanced MDT-15–HIZR-1 binding (Fig 3). This suggests that, besides promoting HIZR-1 nuclear translocation (20), these metals also modulate the TF-Mediator interaction, a classical feature of *bona fide* NHR ligands (54). Importantly, two other HNF4-related NHRs that also bind MDT-15 (34) did not show metal-enhanced binding. This agrees with the notion that zinc is a specific ligand of HIZR-1 (20) and is not involved in the MDT-15–NHR interactions through the zinc-containing DBDs of NHRs (21).

In the Y2H system, we observed increased binding between HIZR-1 and MDT-15 in the low micromolar range of zinc and cadmium. Murine ZIP4 undergoes zinc-stimulated endocytosis when zinc is present in the low micromolar range (55). Similarly, various human and mouse ZIP proteins promote zinc transport in the low micromolar range (56). Thus, the concentration of zinc required to modulate the HIZR-1–MDT-15 interaction resembles that known to engage other zinc homeostasis proteins. This argues that the zinc-modulation of the protein interaction we observed in the Y2H system is likely relevant physiologically.

Other physical interactions are likely involved in this zinc homeostasis mechanism, potentially linking HIZR-1 or MDT-15 to ELT-2 or CDK-8, which we found modestly affects metal regulated genes (Fig 1). In addition, HNF4-like NHRs form both homo and heterodimers (21). It would be interesting to examine whether HIZR-1 forms homo- or heterodimers and whether such interactions are modulated by zinc.

### *mdt-15* protects worms from excess zinc and cadmium

In the *C. elegans* intestine, zinc is stored in lysosome-related organelles called gut granules. In excess zinc conditions, gut granules import surplus zinc via the CDF-2 transporter (44), while zinc is likely also sequestered by MTL-1 and −2. MDT-15 and HIZR-1 promote the induction of these genes when zinc is in excess. This induction is likely physiologically relevant as mutation of either factor results in reduced zinc storage in gut granules and reduced organismal metal tolerance (Fig 4 and (20)).

The study that established zinc as a direct ligand for HIZR-1 did not test whether this NHR also bound cadmium (20). Our data suggest that this may be the case, as cadmium stimulates MDT-15–HIZR-1 binding as strongly as does zinc (Fig 3), and *mdt-15* mutants were sensitive to both metals (Fig 4). However, we note that cadmium could also act indirectly be displacing zinc from other molecular sites and thus making it available for HIZR-1, resulting in higher available zinc levels that indirectly engage the MDT-15–HIZR-1 complex. It will be interesting to define whether HIZR-1 is indeed a direct sensor of the pollutant cadmium.

MDT-15 also promotes oxidative stress responses (36,37,42) and it is conceivable that this function helps protect worms in conditions of excess metals, as e.g. cadmium exposure provokes the formation of reactive oxygen species (7,49). It will be interesting to determine whether HIZR-1 regulates genes other than those involved in storing and mobilizing zinc, such as general or oxidative stress protective genes.

### Mammalian MED15 also regulates heavy metal stress responsive gene transcription

*C. elegans* MDT-15 plays an important role in many adaptive and stress response pathways. Of those, its role in regulating lipid metabolism appears to be conserved in yeast and in mammals (43,57), and its requirement in detoxifying xenobiotic molecules is conserved in fungi (28,58). However, whether mammalian MED15 proteins regulate stress responses was not known. Studying a lung adenocarcinoma cell line that responds to cadmium, we found that MED15 depletion compromises the induction of a metallothionein and that MED15 directly binds the genomic regulatory region of this gene in cadmium enhanced fashion. Similarly, mouse Med15 showed zinc induced binding to the regulatory regions of two genes in MIN6 insulinoma cells; although we did not succeed in effectively depleting Med15 in these cells with transfected siRNAs (not shown), we found that these genes are induced by zinc. Thus, we speculate that the increased binding of Med15 likely upregulates their expression in high zinc. In turn, this likely promotes the protection of MIN6 cells from high zinc (Fig 6). A similar mechanism may protect pancreatic islet β-cells from the transiently high zinc levels these cell experience during insulin exocytosis (52,53).

Currently, we do not know what transcription factor cooperates with mammalian MED15 proteins in metal responsive gene expression. MTF-1 induces cadmium responsive genes in mammals and interacts with Mediator (17), although it is not known which, if any, Mediator subunit directly targets MTF-1. *C. elegans* HIZR-1 is by sequence most closely related to mammalian HNF4a, but functionally and structurally may also resemble mammalian PPARa (21,59). In mouse livers, exposure to the PPARa agonist Wy-14,643 induces *Mt1* and *Mt2* mRNA levels; however, induction is modest (approximately 1.8 fold) and delayed (after 72 hours) (60). Thus, this may well represent an indirect regulatory effect. Additional work is required to determine the mechanisms by which Mediator, and potentially MED15, regulate gene expression in response to high concentrations of metals in mammalian cells and tissues.

In sum, our work highlights the HZA–HIZR-1–MDT-15 regulatory mechanism as a critical transcriptional adaptive mechanism to excess zinc and cadmium in *C. elegans*, with a potentially conserved role for mammalian MED15.

## Materials and Methods

### *C. elegans* transcriptome analysis by microarrays

Microarray transcriptome analysis of the *cdk-8(tm1238)* mutant has been described previously (61). The transcriptome analysis of the *mdt-15(tm2182)* mutant was identical to the analysis of the *cdk-8(tm1238)* mutant using Agilent one-color arrays. We identified a total of 1896 spots with an adjusted P-value of 0.05 or less and a fold-change of >2, representing 798 downregulated and 422 upregulated genes. Microarray data have been deposited in Gene Expression Omnibus. Transcriptome profiles of wild-type worms exposed to 100μM cadmium have been described (62). We determined the overlaps between these datasets and calculated the significance of the overlap as described (36).

### *C. elegans* strains and growth conditions

*C. elegans* strains were cultured using standard techniques as described (63) at 20°C; all strains used here are listed in Table S1. Nematode growth medium (NGM)-lite (0.2% NaCl, 0.4% tryptone, 0.3% KH_2_PO_4_, 0.05% K2HPO_4_) agar plates, supplemented with 5μg/mL cholesterol, were used unless otherwise indicated. *Escherichia coli* OP50 was the food source, except for RNAi, for which we used HT115. Zinc (ZnSO_4_) and cadmium (CdCl_2_) were supplemented in noble agar minimal media (NAMM; (44)) and NGM-lite plates, respectively, at indicated concentrations. For qPCR and phenotype analysis, we synchronized worms by standard sodium hypochlorite treatment and starvation of isolated eggs on unseeded NGM-lite plates; the next day, synchronized L1 larvae were collected, placed on seeded plates at the desired densities, and grown until being harvested at the desired developmental stage, as indicated.

### Gene knockdown by feeding RNAi in *C. elegans*

Knockdown by feeding RNAi was carried out on NGM-lite plates with 25μg/mL carbenicillin, 1mM IPTG, and 12.5μg/mL tetracycline, and seeded twice with the appropriate HT115 RNAi bacteria clone from the Ahringer library (Source BioScience 3318_Cel_RNAi_complete). RNAi clones were Sanger sequenced to confirm insert identity. RNAi negative control was empty vector L4440. RNAi clones are listed in Table S2.

### RNA isolation and quantitative real-time PCR analysis

For *C. elegans*, RNA was extracted from developmentally synchronized worms and prepared for gene expression analysis by real-time PCR analysis as described (61). In all samples, we normalized the expression of the tested genes to the average of three normalization genes: *act-1, tba-1,* and *ubc-2.* For A549 cells, RNA was extracted with RNeasy Mini Kit (Qiagen #74106) according to the manufacturer’s protocol and converted into cDNA for gene expression analysis by real-time PCR analysis as described (Grants et al. 2016). The expression of the tested genes was normalized to the average of three normalization genes: 18S rRNA, GAPDH, and GUSB. We used t-tests (two-tailed, equal variance) or nonparametric tests to calculate the significance of expression changes between conditions, as indicated. Statistical tests were performed based on recommendations by GraphPad Prism 7. qPCR primers were designed with Primer3web (64) and tested on serial cDNA dilutions, as described (61). Primer sequences are listed in Table S3.

### Analysis of the *C. elegans cdr-1* promoter and construction of the *cdr-1p::gfp* promoter reporter

Sequences of regulatory elements involved in the zinc, cadmium, or other detoxification/stress responses were identified from the literature, including ARE, MRE, HSE, DBE, HZA, GATA, and TATA box elements (19,65-74). If available, the equivalent *C. elegans* consensus sequence for each regulatory element was identified in the literature, otherwise the eukaryotic consensus sequence was used. The *cdr-1* promoter (−2853 nucleotides upstream from the predicted transcriptional starting site) was then searched for presence of these candidate elements using SerialCloner 2.5.

The *cdr-1p::gfp* reporter was generated by PCR amplification of the genomic region from 2853 base pairs upstream to 11 base pairs downstream of the *cdr-1* start codon (a G>C mutation at the +3 nucleotide was introduced in the reverse primer to mutate the *cdr-1* start codon) using the primers gtcgacTTTGACGATGACAGAAGAAATG and ggatccTGAATCCAAGATACTTGAGACAGT, followed by BamH1-SalI cloning into pPD95.77 GFP (Addgene #1495) to generate SPD771 (cdr-1p-pPD95.77). Mutant transgenes were generated by site-directed mutagenesis of SPD771 using the Q5 Site-Directed Mutagenesis kit (NEB E0554S) and primers cdr-1p_delGATA1F (CCCTACTTTCccgctgCATTATGTCATCGGG) and cdr-1p_delGATA1R (GTTTCTGTTTCAATTGCAGAATAC) to generate SPD803 (cdr-1p-pPD95.77_DEL_GATA1); cdr-1p_delGATA2F (CCCTACTTTCccgctgCATTATGTCATCGGG) and cdr-1p_delGATA2R (AGAACTGTGTTTTGTGATAAAATTATTG) to generate SPD804 (cdr-1p-pPD95.77_DEL_GATA2); fwd _cdr-1p_delHZA (tcgtggcAATTTTATCACAAAACACAGTTC) and fwd _cdr-1p_delHZA (ccctcaccTCAATTGCAGAAT ACCATTTG) to generate SPD884 (pPD95.77-cdr-1PmutHZA); and fwd_cdr-1p_delDAF16 (ccgtCCAGAAAGCTTAAAATTCAAG) and /rev_cdr-1p_delDAF16 (gtggAACGGAAAAATATAATATGTATATACAC) to generate SPD879 (pPD95.77-cdr-1PΔDBE). All plasmids were verified by Sanger sequencing. Transgenic strains were generated by injecting a mixture of 50ng/μl GFP reporter plasmid, 5 ng/μl *pCFJ90[myo-2p::mCherry],* and 95 ng/μl pPD95.77 empty vector into wild-type worms, and then selecting transgenic mCherry-positive progeny.

### Staining of *C. elegans* gut granules with FluoZin-3

FluoZin-3 acetoxymethyl ester (Molecular Probes F24195) was reconstituted in dimethylsulfoxide (DMSO) to a 1mM stock solution. FluoZin-3 was diluted in M9 to generate a concentration of 30μM and dispensed on NAMM plates, as described (44). Synchronized wild-type and mutant L4 stage worms were transferred from NGM-lite plates to these plates and cultured for 16 hrs. Worms were then transferred to NGM-lite plates without FluoZin-3 for 30 min to reduce excess FluoZin-3 signal from the intestinal lumen before imaging.

### Fluorescence microscopy on *C. elegans*

For imaging, worms were transferred onto 2% (w/v) agarose pads containing 15μM sodium azide (NaN3). For the analysis of worms bearing the *cdr-1p::gfp* reporter, the worms were imaged using differential interference contrast (DIC) and fluorescence optics through an HQ camera (Photometrics, Tucson, AZ, USA) on a Zeiss Axioplan 2 microscope (Carl Zeiss Microscopy, Thornwood, NY, USA). Analysis of fluorescence intensity was performed using ImageJ software, normalizing for area and background fluorescence.

For analysis of FluoZin-3 stained worms, we used a Leica SP8 confocal microscope with Leica LAS X software. Images of worms with different genotypes were taken with the same exposure times. To assess zinc storage, we quantified the number of FluoZin-3 stained granules in the first six cells of the gut with ImageJ2 (75). To eliminate non-specific fluorescence, we manually removed background signal outside of the gut cells and in the gut lumen. Background fluorescence in the remaining part of the image was then subtracted equally from all images, and images were smoothed with the “Sigma Filter Plus” (edge-preserving noise reduction) function. Images were simultaneously adjusted with Auto threshold of “MaxEntropy” and Auto local threshold of the mean. After adjusting the threshold and reducing background noise, granules where counted automatically with the “3D Objects Counter” function. To verify the automatic count, we manually counted the number of granules in randomly sampled images.

### *C. elegans* egg laying assay

N2 and *mdt-15(tm2182)* worms were grown from late L4 stage for 24 hours on agar A plates seeded with OP50 and supplemented with 100μM zinc. Eggs and L1 progeny were counted after that time and compared to worms grown on agar A plats with no additional treatment.

### Yeast-two-hybrid assays and Western blots

MDT-15 bait plasmids and NHR-49 and SKN-1c prey plasmids have been described (42). The wild-type HIZR-1 cDNA sequence was amplified with the primers fwd_BamHI_NHR-33_preyY2H (ggatccATGCAAAAAGTTATGAATGATCCTG) and rev_XhoI_NHR-33_preyY2H (ctcgagATCATTTTTCGTATGAACAATGCAC), and cloned into the BamH1 and SalI sites of pGADnewMCS to generate SPD885 (pGADnewMCS_NHR-33). We then used the NEB Q5 site-directed mutagenesis kit (E0554S), template SPD885, and primers SDM_nhr-33-am285_F (AGCAGAAaATGCTGCAAAAAT) and SDM_nhr-33-am285_R (GATTGTACACCCTCTGCATTC) to generate SPD918 (pGADnewMCS_NHR-33_am285). All plasmids were sequenced to verify the accuracy of the sequence amplified by PCR and the absence of other mutations. Pairs of plasmids were transformed into strain Y187 (Clontech, Mountain View, CA, USA) and liquid β-galactosidase assays were performed using an OMEGASTAR plate reader (BMG Labtech, Ortenberg, Germany), as described (42). Each assay included at least three technical replicates and was repeated three or more times. Yeast lysis, SDS-PAGE, and Western blots to detect protein expression were done as described (42). Antibodies used were GAL4 AD Mouse Monoclonal Antibody (Takara Bio USA, Inc. #630402), GAL4 DNA-BD Mouse Monoclonal Antibody (Takara Bio USA, Inc. #630403), and GAPDH Mouse Monoclonal Antibody (CB1001-500UG 6C5) for normalization.

### Mammalian cell culture and transfection

A549 lung adenocarcinoma cells were obtained from ATCC and maintained in Dulbecco’s Modified Eagle Medium (DMEM; Gibco #11995065) supplemented with 10% fetal bovine serum (FBS; Gibco #12484028), as described (51). The cells were seeded at a density of 5×10^4^ cells per well in 24-well plates 24 hours before transfection. On the day of transfection, scrambled (Dharmacon #D-001206-13-05) and MED15 specific (Dharmacon #M-017015-02-0005) siRNAs were delivered into A549 cells using DharmaFECT 1 Transfection Reagent (Dharmacon #T-2001-03) according to the manufacturer’s protocol. The transfection medium was replaced with complete medium after 24 hours, and the cells were treated with 5μM CdCl2 for hours 48 hours post-transfection. MIN6 cells were cultured in 25mM glucose DMEM, as described (76), and zinc stimulation was performed by addition of 50μM ZnSO4 for 24 hours.

### ChIP in A549 and MIN6 cells

ChIP was performed as described (77) in A549 and MIN6 cells with minor modifications. Briefly, cells were grown for 24 hours with or without 50μM CdCl2 and 100μM ZnSO4 (respectively) to 2×10^7^ cells per plate, and crosslinked by adding paraformaldehyde to a final concentration of 3% for 10 minutes at room temperature. Immunoprecipitation was performed using MED15 antibody (ProteinTech, 11566-1-AP). Crosslinking was then reversed by overnight incubation at 65°C and DNA purified using QIAquick PCR purification column (Qiagen, 28104). Immunoprecipitated DNA was then quantified via Qubit (ThermoFisher, Q32854) and analyzed by qPCR for appropriate genomic regulatory loci and controls. Primer sequences are listed in Table S3.

## Acknowledgements

We thank the Taubert lab and Dr. Frédéric Picard (Université Laval) for comments on the manuscript. Some strains were provided by the CGC, which is funded by NIH Office of Research Infrastructure Programs (P40 OD010440). We thank Rocky Shi (BCCHR) for helping us develop the script to quantify gut granule number in the *C. elegans* intestine. Grant support was from The Canadian Institutes of Health Research (CIHR; PJT-153199 to ST), the Natural Sciences and Engineering Research Council of Canada (NSERC; RGPIN-2018-05133 to ST), the Cancer Research Society (CRS; to ST), and Diabetes Canada (OG-3-15-4946-FL to FL and ST). NS was supported by NSERC CGS-M and UBC Medical Genetics scholarships, AZK by a BCCHR Canucks for Kids Diabetes laboratory scholarship, JMG by a Vanier Canada Graduate Scholarship, MYYL by a BCCHR summer studentship, AM by an NSERC-USRA studentship, and ST by a Canada Research Chair. No author has any conflict of interest.

## Supporting Information

Table S1: List of worm strains.

Table S2: List of HT115 RNAi bacteria clones from the Ahringer library.

Table S3: List of primers used in qPCR experiments.

Figure S1: The *cdr-1p::gfp* transcriptional reporter is induced by 100μM cadmium and 200μM zinc, and induction in adult worms by cadmium requires *elt-2.*

Figure S2: MDT-15 and HIZR-1 interact physically.

Figure S3: *mdt-15(tm2182)* and *hizr-1(am286)* mutants have a zinc storage defect.

Figure S4: Expression analysis of fusion proteins for Y2H assays.

